# In pursuit of visual attention: SSVEP frequency-tagging moving targets

**DOI:** 10.1101/804708

**Authors:** Peter de Lissa, Roberto Caldara, Victoria Nicholls, Sebastien Miellet

## Abstract

Previous research has shown that visual attention does not always exactly follow gaze direction, leading to the concepts of overt and covert attention. However, it is not yet clear how such covert shifts of visual attention to peripheral regions impact the processing of the targets we directly foveate as they move in our visual field. The current study utilised the co-registration of eye-position and EEG recordings while participants tracked moving targets that were embedded with a 30 Hz frequency tag in a Steady State Visually Evoked Potentials (SSVEP) paradigm. When the task required attention to be divided between the moving target (overt attention) and a peripheral region where a second target might appear (covert attention), the SSVEPs elicited by the tracked target at the 30 Hz frequency band were significantly lower than when participants did not have to covertly monitor for a second target. Our findings suggest that neural responses of overt attention are reduced when attention is divided between covert and overt areas. This neural evidence is in line with theoretical accounts describing attention as a pool of finite resources, such as the perceptual load theory. Altogether, these results have practical implications for many real-world situations where covert shifts of attention may reduce visual processing of objects even when they are directly being tracked with the eyes.

## Introduction

In daily life we experience a large variety of situations in which we need to visually track multiple objects at the same time, for instance when crossing a busy street, monitoring the safety of children playing in playgrounds, locating an errant spouse in a bustling shopping centre, etc. In these situations we rely on the division of visual attention as we monitor both moving and stationary objects across time, often rapidly switching between attending to targets through direct eye-movements or through our peripheral visual fields. The need to modulate our attention arises from inherent limitations in our capacity to attend to the broad array of stimuli our senses may provide to us at any one moment [1-2]. The Perceptual Load Theory advanced by Lavie and others conceptualises attention as a limited pool of resources that we are able to devote to the processing of targets and distractors in various environments. The balance of our attention directed to spatial locations at any given moment is thus related to the perceptual load of the tasks being concurrently performed [3-6]. The way in which the brain modulates visual input through attention has additionally been conceptualised as a mechanism that decreases the salience of distractors by reducing the neural sensitivity to unattended stimuli so that attended stimuli experience less competition while they are processed [7-9]. When applied to contexts and tasks that require the visual analysis of complex scenes involving both moving and stationary objects, these theories [3-6] suggest a modulation of attention that depends on the requirement of attention to be either divided or singularly focused. Part of such a dynamic involves attention directed to what we are directly foveating on (overt attention), as well as attention directed to areas outside our foveal fields in our parafoveal or peripheral visual fields in the form of covert attention [10]. While overt visual attention can be indexed through the recording of eye-position during various tasks, covert shifts of attention to areas outside of foveal regions are by their nature often not accompanied by explicit behavioural measures and must be measured indirectly through analyses of reaction time in paradigms involving cueing to extra-foveal spatial locations compared to either an un-cued or an incorrectly cued location [11-12].

Aligning with the view of attention to be a limited pool of resources, some studies have suggested that when covert attention is directed to spatial areas in the periphery there is a decrease in attention directed towards foveated stimuli [13-14]. However, competing evidence has suggested that both covert and overt visual attention may be deployed simultaneously in parallel in paradigms involving dual tasks without a notable decrease in performance [15-16]. The nature of the tasks in such paradigms is likely to play a critical role in how attention might be divided between overt and covert monitoring during analysis of objects in the visual environment. In the case of complex scenes this may involve a selection of what targets to monitor overtly with the eyes and which to monitor covertly through the shift of peripheral visual attention. What is yet to be clarified is how overt attention directed to a moving object is influenced by additional requirements to monitor other spatial locations with covert visual attention. This question is the basis of the current study.

While various behavioural tasks have been used to investigate the deployment of both overt and covert visual attention, it is possible to index the relative recruitment of these forms of attention by recording the neural responses in electroencephalographic (EEG) recordings to the flickering of stimuli presented in different spatial locations of the visual field. While the early occipital lobe responds to this flickering in a systematic way, the strength of this response is strongly modulated by whether the flickering objects/regions are being attended to or not, with larger responses to attended stimuli compared to unattended stimuli [17-22]. Known as Steady-State Visual Evoked Potentials (SSVEP), this technique offers a complement to the measurement of behavioural responses, as it can capture the time-course of shifts of attention, contrasting with behavioural responses which, while influenced by attention, constitute the end-point of a chain of perceptual and decision-making processes. Indeed, an important benefit of the SSVEP approach is that it does not require a specific behavioural response, making it well-suited to investigate shifts of attention that take place without behavioural markers [23]. Recent studies investigating attention allocation during smooth-pursuit paradigms have found clear neural responses to flickering stimuli in both peripheral regions [24] and to a general flickering background stimulus [25], with the latter suggesting the neural responses during smooth-pursuit to be larger than when the eye-position is fixed. However, to our knowledge this paradigm has not yet been used to investigate overt visual attention during the tracking of moving objects or how it is affected by task-related shifts of covert attention.

Applied to the question of how visual attention is affected by the need to attend to peripheral areas while simultaneously tracking a moving object, the SSVEP technique offers a means of determining whether such covert shifts of attention decrease the sensitivity to the moving foveal target as might be predicted if a limited pool of visual attention leads to a sacrifice of overt visual attention when deploying covert attention. In order to investigate this question while maintaining systematic control over low-level visual properties, the current study combined eye-position recordings with an SSVEP paradigm, measuring the neural responses to the flickering of targets as participants followed them with their eyes as they moved across a computer screen. The task consisted of overtly tracking a target as it moved across a computer screen and pressing a button when it entered a specific portion of the screen. However, in half of the trials the participants were instructed that a second target might also appear and follow the same trajectory as the first, whereupon they should perform the task on the second target instead. This manipulation created two conditions: An undivided condition where the task required overt attention only to one single moving target, and a divided condition where the expectation of a possible second target at a specific place and time provided the context for both overt and covert visual attention to be used in the task. The neural responses to the foveally-tracked target thus formed an index of overt attention, which would be significantly reduced in the case of shared and limited pool of attentional resources, when the participants covertly monitored for the appearance of a second target.

## Methods

The Human Ethics Committee at the University of Fribourg approved the methods and procedure used in this study.

### Participants

22 participants were tested in the current study. 4 participant datasets were excluded due to insufficient trial numbers to form a meaningful condition average and one dataset was excluded due to strong contamination throughout the scalp originating from anterior/facial areas during the trials which introduced distinct 10 Hz distortions and a 20 Hz harmonic [26-27]. Datasets were analysed from the remaining 17 participants (13 females, 17 right-handed), aged between 19 and 44 years (mean age = 26.5 years, SD = 7). All participants had normal or corrected-to-normal vision, and gave their informed consent before participating in the study. Participants were offered 50 CHF for their time or course participation credits.

### Stimuli and procedure

Participants were instructed to follow a moving target as it moved across a computer screen and to press a keyboard button when the target entered a spherical “goal” portion of the screen. The targets consistently travelled along a diagonal path from the top-left part of the screen to bottom-right goal section (see Fig 1) at a speed of 3.75 deg/s. The target stimuli consisted of a black and white rectangle (1.05° x 2.10° visual angle) checkerboard pattern alternating (reversing between black and white) at a consistent rate of 30 Hz against a white background. The 30 Hz flicker created the frequency tag used for the subsequent EEG analysis of visual attention.

**Fig 1.**
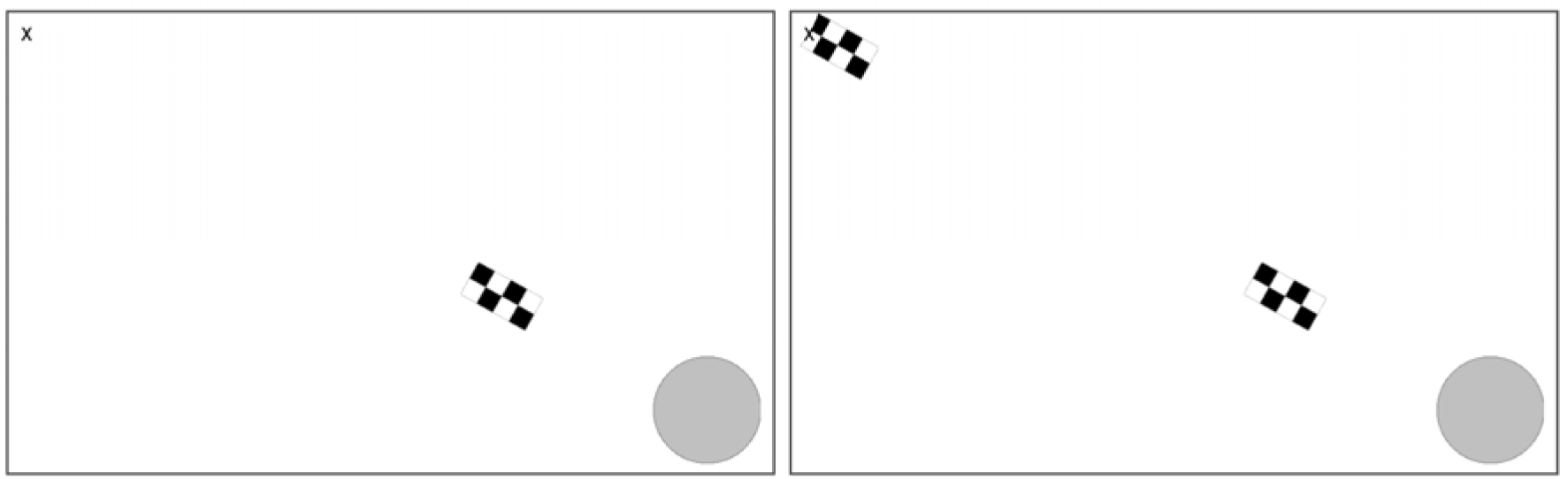
Undivided and divided attention trial examples. Alternating checkerboard targets emerged from the left side of the screen (denoted by “x”) and travelled across the screen to a circular region (left), whereupon participants pressed a button when they judged the target to be fully within the region. In half of the trials a second target had a 66% probability of appearing and travelling across the screen (right), where participants had to subsequently perform the button-press task on this second target instead.

Two experimental conditions were created by manipulating what participants expected to see in the trials. In one condition block, participants were instructed that only one target would travel across the screen in each trial, and that they should press the keyboard button when it reached the goal area. In the other condition block, participants were instructed that a second target might appear while the first target was still travelling across the screen (occurring in 2/3 of the trials in this condition). The second target (also flickering at the 30 Hz) appeared in the same location as the first target at the onset of the trial so that participants had a specific and predictable target for a covert shift of attention. The second target appeared at either 2136 or 2270 ms into the trial, providing a consistent time range for participants to predict when to direct covert attention with a small range of variability. Participants were instructed that if a second target did appear they were to then track that second target with their eyes and press the button when the second target reached the goal area. This was to provide a task-related division of attention, while balancing all low-level visual properties between the conditions up until the appearance of a second target in the periphery. Participants were informed at the beginning of each condition block whether to expect either only one or more than one target, creating two experimental conditions; an undivided attention condition and a divided attention condition. The time-window leading up to the possible presentation of a second target thus formed the period of interest for our analysis, where shifts of attention relating to participants condition-related expectations were predicted to occur. Analyses were not performed on the time-range beyond this period of interest, as the conditions following this time diverted in both low-level properties (competing foveal and peripheral flickering stimuli) as well as in the behaviours of the participants (continued tracking of single target versus saccadic responses of varied latency to the second target).

There were 204 trials in total, with 102 in each of the divided and undivided attention conditions. The trials were divided into 4 alternating homogenous blocks, and the presentation order of these blocks was counter-balanced to avoid fatigue or order-effects by creating two block-orders presented to two participant groups (8 and 9 participants in the two counterbalanced groups). The experimental trials began with a fixation cross in the top left corner of the screen, corresponding to the region where the target would initially appear. When participants fixated on this cross area (1.3° x 1.3°), the cross would disappear and the target would begin to emerge from the top-left corner 266 ms later. The trials ended when the participant made their decisions relating to the targets completely entering the goal area (3170 ms into the trial in the undivided condition, 5003 or 5136 ms into the trial in the divided condition) by making a keypress. The experimental stimuli were presented on a 24 inch VIEWPixx/3D monitor (1920 x 1080 pixels, 120 Hz refresh rate) at a distance of 75 cm, and presented through Experiment Builder (v1.10.1630) software.

### Eye-movement recording and processing

Eye-positions were recorded through a desktop-mounted Eyelink 1000 monocular (left) eye-tracker sampling at 1000 Hz. Calibrations of the eye-tracker (13-points, average position error < 0.5°) were performed at the beginning of the experimental blocks and after breaks in the trials. The onset of a trial was triggered by a fixation in a specified region in the top left part of the screen; if this was not fixated upon within 4 seconds after presentation then a re-calibration sequence was entered, ensuring effective calibration throughout each of the trials. The trials began with the flickering targets emerging near the upper-left portion of the screen 266 ms after trial onset. The early part of the trials was characterised by the target stimuli approaching and passing the participants’ fixated gaze, and the subsequent orienting of their gaze to these moving targets through catch-up saccades. This orienting phase generally took approximately 500 ms before participants were able to align their smooth-pursuit eye movements with the movement of the targets. To allow for this, a time-window of analysis for eye-gaze and EEG was created, beginning 1000 ms after the onset of the trial (700 ms after the onset the first target) and ending at 2000 ms (shortly before a second target might appear at either 2136 or 2270 ms).

The x and y gaze coordinates of the participants in the trials were exported and analysed to ensure that the flashing targets were directly foveated by the participants during a 1000 ms period immediately preceding the time at which the onset of a second target would occur. Trials were rejected if the participants were not directly foveating the targets for over 95% of this 1000 ms time period (allowing for transient loss of foveation and eye-blinks). A 1000 ms period of interest was chosen for two reasons: 1. It is preceding the likely appearance of the second target so we expect relevant processes associated with attentional shifts to occur in this period, and 2. This period starts after the catch-up saccade and when the smooth-pursuit is consistently initiated across trials. To ensure a reliable average, a critical threshold of 25 accepted trials was applied, which led to the rejection of 4 participants due to insufficient trials. The SSVEP technique has been found to yield a high signal to noise ratio, with analyses involving known oscillations (frequency tags) reliably measuring visually-entrained EEG responses from as little as 10 artefact-free trials [28], and from 15 trials in a face-detection paradigm using sweep SSVEP [29]. Because the co-registration of EEG and eye-movements in the current study required rejection of EEG epochs where eye-gaze was outside of the stimulus regions, the trial rejection rate was considered in the experimental planning, with 102 trials per condition being chosen to allow for a large number of rejected trials, equating to an average rejection of 39% of all trials. The average number of accepted trials in the divided and undivided attention conditions in the current study was much higher than this minimum threshold, with 65 and 60 accepted trials, respectively (see supporting Fig 1). Statistical analysis conducted on the accepted trial means for each condition showed no significant effect (*t*(16) = 1.71, *p* = .108, *d* = 0.41). This analysis was complemented with a Bayes factor analysis, which allows for an interpretation of not only the likelihood of the data representing evidence in favour of a hypothesised difference between conditions, but of evidence in favour of a null-effect [30]. The results suggested anecdotal evidence [for review of Bayes factor see 31] for the null hypothesis and no support for predicted differences between the conditions (*BF*_*10*_ = 0.825, 0.005% error). Bayes factor and t-test analyses were performed using JASP (0.11.1) with default settings [32].

After the trial exclusion process, the remaining trials were analysed to determine whether there were systematic differences in eye-position between the divided and undivided conditions. Repeated measures two-tailed permutation (100,000 permutations) *t*-tests using a *t*max statistic [33] were performed at each time point in the period of interest for the X and Y eye-position data, which did not reveal significant patterns of difference between the conditions. This was also performed on data that indexed the absolute distance (in degrees of visual angle) between the participants’ eye-positions and the centre of the target at each time point, which similarly yielded no distinct pattern of difference (the most extreme *t*/*p* during the critical period was *t*(16) = −1.57, *p* = .834). To supplement the *t*-test analyses, Bayes factor analyses were additionally performed on the absolute distance data at each time point across the period of interest to determine the likelihood that the means were indeed comparable. The results indicated greater support for the null hypothesis (divided absolute distance = undivided absolute distance) than a genuine difference across this time period, with an average *BF*_*10*_ of 0.307 and a maximum of 0.566 (see Fig 2c for *BF*_*10*_ values across the period of interest). The participants’ accuracy at tracking the targets can be seen in Figs 2a, 2b, and 2c, which depicts the distances at each time point that the participants’ eyes were from the target centre in XY co-ordinates and in absolute Euclidean distance, measured in degrees of visual angle. A value of zero would therefore correspond to the centre of the target in either the X or Y plane. The high target-tracking accuracy in the current study is consistent with previous studies utilising targets of predictable speeds [34], and is in line with the results of a previous study showing that following the centre of a moving target facilitates the allocation of attention to peripheral locations when multiple objects are present [35].

**Fig 2.**
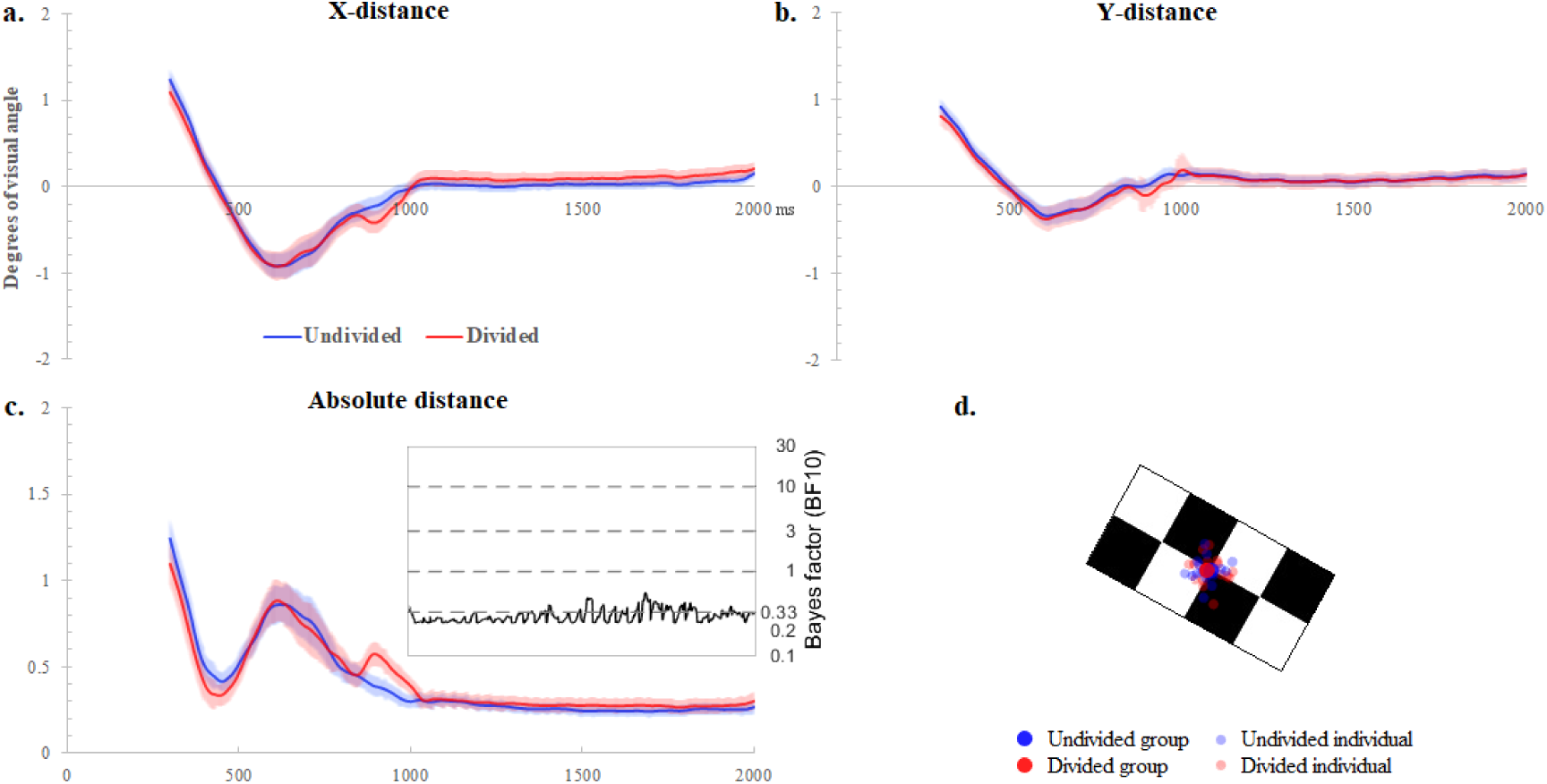
Eye-gaze accuracy during target tracking. Eye-gaze distance from target centre (at 0, measured in degrees of visual angle) for x (a), y (b), and absolute (c) co-ordinate dimensions from the onset of the target until immediately before the possible appearance of a second target (standard error shaded). Bayes factor analyses performed on absolute distance did not suggest a reliable difference between the two conditions (Fig 2c). Fig 2d depicts the condition average eye-positions (solid large blue/red) relative to the targets throughout the 1000 ms interest period, as well as the individual participant averages (faded small blue/red).

The average distance from target centre for the undivided and divided attention conditions were −0.05° and 0.01° respectively for the X positions, and 0.03° and 0.02° for Y positions. For reference, the length and width of the target stimuli were 2.2° and 1.1° respectively. The average precision (both group average and individual average) across the critical time-period is illustrated in Fig 2d, which represents the average eye-positions on the target throughout the period of interest for the divided and undivided conditions.

To determine whether the incidence of eye blinks or saccades in the current study was modulated by condition, the mean number of blinks and saccades each participant exhibited in each critical period per condition was analysed through repeated measures paired *t*-tests and Bayes factor analysis. Blink and saccade events were detected through Dataviewer (version 1.11.900) using saccadic velocity and acceleration thresholds of 30°/sec and 8000° /sec^2^, respectively. The results indicated no significant differences in the incidence of eye blinks or saccades between the conditions (*t*(16) = 0.868, *p* = .398, *d* = 0.21; *t*(16) = −1.438, *p* = .17, *d* = −0.39), and Bayes factors (*BF*_*10*_) of 0.346 (0.003% error) and 0.595 (0.002% error) respectively, indicating anecdotal evidence in favour of the null hypothesis (no difference between conditions). On average the number of blinks per trial were very low, with 0.25 (SD=0.26) blinks per trial in the undivided condition and 0.22 (SD=0.23) in the divided condition, as was the number of saccades (1.07, SD=0.59, and 0.99, SD=0.51, respectively).

Participants’ eye-movements were monitored during the testing session by the experimenters to ensure they understood and followed the task instructions. Participants initiated a saccade to the second target within 351 ms (SD=70 ms) of their appearance in relevant trials, suggesting an adherence to the task instructions. Following the pre-processing of eye-position data, only accepted trials were used in subsequent statistical analyses of task-related effects on EEG responses.

### EEG recording and processing

Electrophysiological responses were recorded through a Biosemi Active-Two amplifier system, using 128 Ag/AgCl electrodes sampling at 1024 Hz. Additional electrodes were placed at the outer canthi and above of each eye, to register ocular movements and blinks. EEG data was processed offline through EEGLAB (14.1.0b) running in the MATLAB 2016B environment. After an initial bandpass filtering process (0.1-75 Hz, zero phase shift, linear finite impulse, Hamming window), epochs of 5000 ms duration were created, beginning at a −1000ms baseline period at the onset of the trial. To isolate and remove blink and eye-movement distortions, the 5000 ms epochs were subjected to Independent Component Analysis (ICA, using the ‘runica’ algorithm through EEGLAB) [36]. Independent components corresponding to frontal blink and saccade topographic distortions were isolated and removed from the data, as well as slow drift in EEG corresponding to smooth pursuit activity (see supporting Fig 2). However, in a number of datasets this slow drift was not able to be isolated through ICA, even though a clear drift could be observed in the raw data. This was not problematic in the current experimental design, however, as the slow drift was not related to frequencies overlapping the 30 Hz frequency tag utilised in the study (see supporting Fig 2a and 2b for examples of the frequency responses of independent components associated with blink and smooth pursuit distortions).

The EEG was subsequently re-referenced to a common-average reference, and epochs noted for rejection in the eye-gaze analysis were removed from statistical analysis, leaving only epochs where the participants were directly foveating the targets more than 95% of the critical 1000 ms period. Frequency power values were measured relative to a 1000 ms pre-stimulus onset baseline to quantify event-related spectral perturbation (ERSP) data in a normalized signal-to-noise ratio (SNR), and are hereafter presented in dB units (relative to pre-stimulus baseline). The frequency tag from a directly foveated flickering stimulus was predicted to lead to a corresponding neural frequency in the central-occipital region [37], approximately between central Oz and Iz electrodes in a 10-20 system. This was confirmed with a fast-fourier transform of the full 1000 ms critical period, where a 30 Hz signal was observed in the central occipital region relative to the 1000 ms baseline period (see Fig 3a for a topographical representation of 30 Hz power). The frequency response spectrum at the posterior occipital cluster indicated a discrete spike in the 30 Hz frequency band (Fig 3b). This was complemented with a time-frequency decomposition using Morlet wavelet transformations within the range of 3-70 Hz (3 0.5 wavelet cycles; yielding higher resolution as frequency increased and a wavelet at exactly 30 Hz) to give insight into the timing of the 30 Hz signal from the beginning of the trial to the period immediately preceding the possible onset of a second target (2000 ms window), collapsing across the two conditions. The 30 Hz signal was observed in both the ERSP and inter-trial coherence (ITC) topographies to arise at approximately 750 ms in the central occipital region and continuing through to the end of the 2000 ms window. Differences in 30 Hz power preceding the beginning of the 1000 ms period of interest were not systematically balanced for low-level differences in visual perception, as the gaze of the participants was not ensured to be on the flickering targets at this time, and thus not analysed.

**Fig 3.**
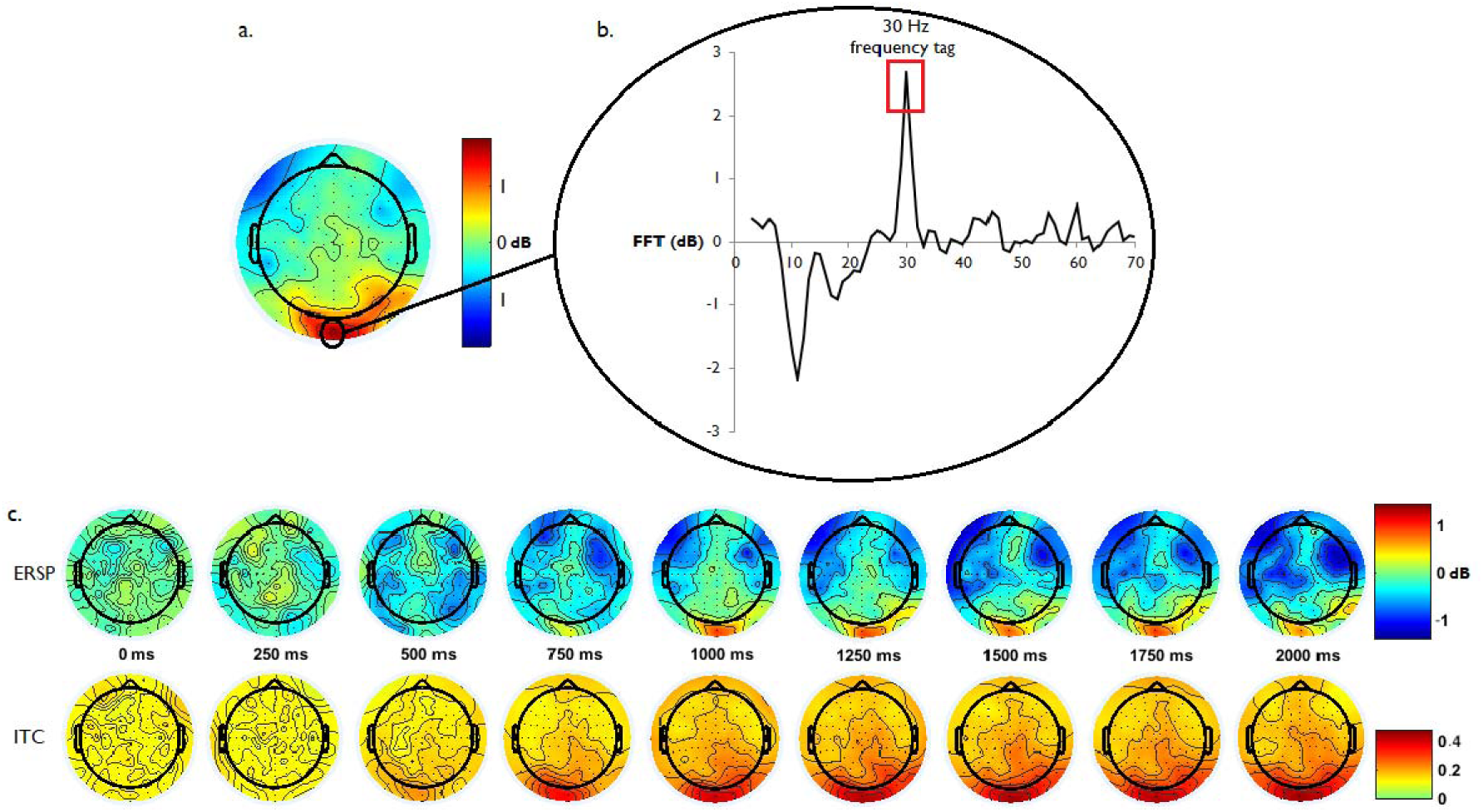
Scalp topography of 30 Hz SSVEP signal. Scalp topography revealed a strong 30 Hz signal in the central-occipital region during the 1000 ms critical period (a), with a fast-fourier transform in this area indicating a distinct 30 Hz spike corresponding to the frequency-tag (b). Event-related spectral perturbation (c) and inter-trial coherence transforms found reliable 30 Hz signatures in the central-occipital regions arising at approximately 750 ms into the trial and continuing through the target-tracking period.

Following the confirmation of the 30 Hz frequency tag in the EEG recordings, statistical tests were conducted to compare the effect of divided visual attention on the power of the mean oscillation in the midline posterior occipital region corresponding to the Oz and Iz electrodes for all participants, representing the two electrodes with the largest 30 Hz signals (Fig 3a). Event-related spectral perturbations (ERSP) from the Morlet wavelet transformations from this region were computed for the divided and undivided attention conditions, producing ERSP averages of each condition for each participant. Differences between the divided and undivided ERSP data at each time point in the 1000 ms period of interest were compared with both Bayes factor analyses, and a repeated measures, two-tailed permutation (100,000 permutations) test using the *t*max statistic implemented through R [33]. The Bayes factor analysis gave an index of whether the 30 Hz ERSP power data provided support for hypothesised differences between the conditions across the time range, or whether a null-effect was more likely. The non-parametric *t*max procedure provided a statistical index of the reliability and direction of any observed differences in the ERSP data, corrected for multiple time-point comparisons by permuting the observed 30 Hz ERSP data to arrive at adjusted *t* and *p*-values.

## Results

Bayes factor analyses of the difference in 30 Hz power between the divided and undivided conditions (Fig 4 a & b) in the period of interest showed substantial evidence for hypothesised differences early in the time window (*BF*_*10*_ > 3) from 1289 through to 1367 ms, and strong evidence from 1317 to 1331 ms (see Fig 4c). Of note are results of the Bayes factor analyses for the other time points in the period of interest, where the *BF*_*10*_ values suggest substantial support for the null hypothesis, or no differences between the divided and undivided conditions (*BF*_*10*_ < 0.33). The non-parametric *t*max analyses also revealed a period of significant difference (*p* < 0.05) in the same early time-range, arising between 1307 ms and 1343 ms after 1^st^ target onset, corrected for multiple comparisons. The source of these differences was observed to be due to greater 30 Hz power in the undivided attention condition compared to the divided attention condition. This difference can be observed in Fig 4c and 5. The shaded permutation-corrected 95% confidence intervals in the difference wave in Fig 5 suggests that this was a modest effect.

**Fig 4.**
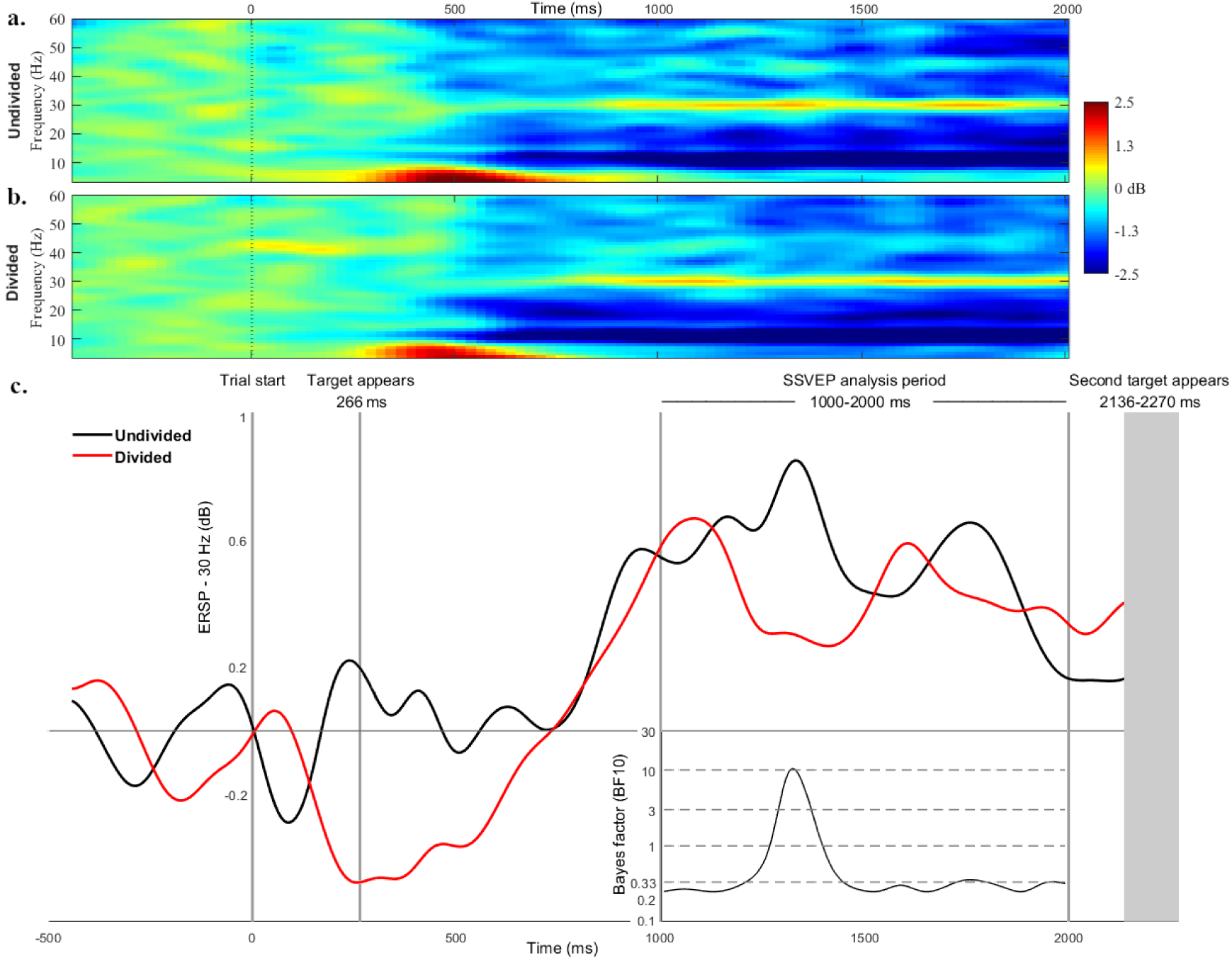
30 Hz SSVEP power. 30 Hz power (ERSP) across the trial period indicated sustained attention in both the undivided condition and divided attention conditions (4a & 4b), with apparent decreases in neural response at discrete periods in the divided attention condition (4c). The results of Bayes factor analyses suggested an early discrete period of difference between the conditions, while all other time points the 30 Hz signal were not different. Grey lines denote the timing of events in the trials.

**Fig 5.**
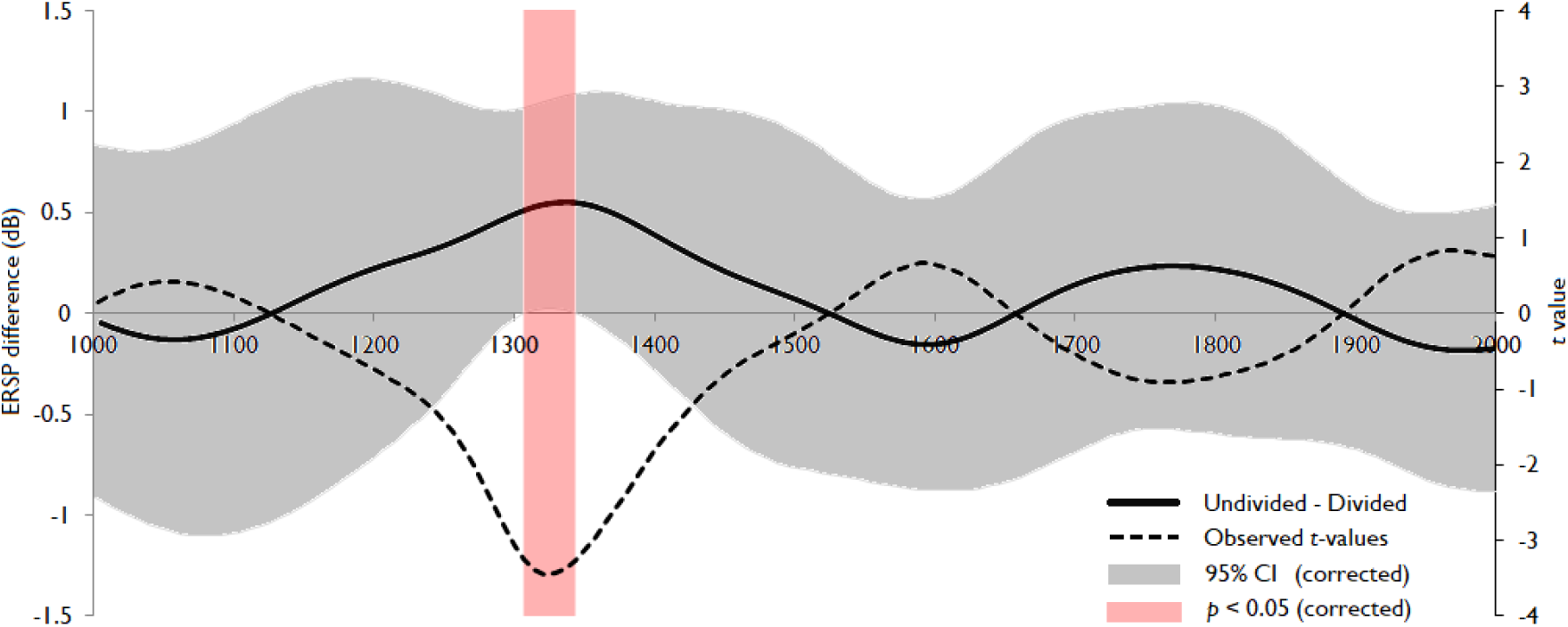
*T*max permutation analysis results. Analyses of differences in the power of the 30 Hz frequency tag in the undivided and divided attention conditions over the 1 second period of interest revealed significantly lower 30 Hz power in the divided condition at approximately 1300 ms (precise window shaded red). 95% confidence intervals in the subtraction waveform (undivided minus divided) denote periods of significance (p < 0.05) when they do not overlap zero.

## Discussion

The current study sought to measure overt visual attention in a smooth-pursuit paradigm, and to determine whether allocation of covert attention to peripheral regions modulated measures of overt attention to the moving target. The SSVEP power corresponding to the 30 Hz frequency tag of the moving stimuli decreased when the task required participants to attend covertly to where an additional target might appear in the periphery while concurrently tracking a moving target. This finding aligns with the view of covert and overt visual attention as expressions of a pool of attentional resources, where an increase in covert attention can lead to a concomitant reduction in overt attention [1-2], similar in nature to the reduction in SSVEP power to foveated static stimuli observed when covert visual attention is recruited [13]. Our findings indicate that attention can be deployed covertly while tracking a moving target, which is also in line with the behavioural results of [38], who used saccadic response times to index covert attentional shifts to peripheral spatial regions. Similarly, they support the behavioural findings of Ludwig et al. [16] suggesting that both covert and overt attention can operate in parallel. A notable difference between our methodology and that of Ludwig et al., however, is that we utilised a passive measure of overt visual attention through SSVEP rather than a behavioural index. Thus, our approach allowed us to investigate the fine-grained temporal modulations of overt attention resulting from allocation of covert attention, rather than the end-product. While our SSVEP results indicate that there was a reduction in overt attention to the moving targets when the task required a covert shift of attention to a peripheral location, this, however, does not necessarily mean that a performance decrease would also be observed had an additional behavioural task been employed. This is in line with the perceptual load theory, which suggests that the division of visual attention across covert and overt areas is moderated by the processing load required by the tasks at hand. Accordingly, it is likely that modulating the salience of the moving target may also modulate the degree to which covert shifts of attention to peripheral locations affect the processing of the moving target. In contexts such as parents tracking moving children in a playground, or security forces monitoring moving threats, one dimension of the task involves accurately following targets with the eyes while an additional task might involve a specific visual analysis of the target itself. In these contexts, the level of overt attention may be higher than when there was no secondary task requiring visual analysis, making it more difficult (or less likely) for covert shifts to occur. It is also likely that additional visual analysis of the moving stimuli would require greater overt attention and thus may limit the amount of covert attention available for monitoring other spatial areas, as suggested by the finding that foveal distractors are harder to ignore than peripheral distractors [39]. The nature of any such task will likely then influence the relative strength of both central overt and peripheral covert visual attention, as competition between features for visual analysis and their distractors in central vision has been found to lead to an enhancement of neural sensitivity to peripheral regions [40].

However, it is important to note the transience of the period of reduced neural sensitivity to the moving targets in the current study, especially considering that this period of significant difference was relatively early in the trial. The current study utilized expectation of the likely appearance of a second target in the trials to create a task-related division of visual attention between a moving target and a defined area in the participants’ left peripheral field. While this allowed the participants to know where to allocate covert attention in these trials, the timing of the appearance of second targets was also somewhat predictable. While it might have been a logical prediction that the effects of divided attention in these trials would be more likely to be observed as time advanced towards this critical moment, our data did not show this. Rather, differences in the neural response to the overtly tracked targets were reliably observed approximately 800 ms before the time a second target would have appeared. The results of the Bayes factor analysis (Fig 4c) provides a useful insight into the timing of the effect, and reveals that if anything there was anecdotal evidence that the conditions were comparable at other times, rather than merely failing to be significantly different. It is possible that within any one trial there are multiple discrete shifts of attention away from the moving targets, but that the timing or duration of such shifts are not systematic within the trials and thus do not reveal a statistically clear pattern when averaging across the them. Such an interpretation also invites speculation as to why the period or reliable difference in SSVEP power occurred early in the trial. Such an early, discrete period of reduced overt attention may reflect a process of the encoding of spatial locations for future monitoring through covert attention, where overt attention is impacted to a lesser degree after this encoding process has occurred. The reduction in SSVEP power during this early period could then be considered to be due to processes involving the future execution of eye-movements, rather than an ongoing sampling of covert visual areas in order to react quickly to a second target appearing. If the predictability of the time or location of potential distractor stimuli modulates the time or the strength of changes in overt visual attention during object tracking, then future studies might specifically manipulate these dimensions to determine how they contribute to such effects, and whether they interact with the task requirements.

Additionally, there are other low-level factors that might be predicted to modulate both overt and covert visual attention during smooth-pursuit, such as the speed of the moving target, and the spatial locations of where covert shifts of attention are directed. Saccade latencies to stimuli presented during smooth-pursuit have been found to increase as target speed increases [38, 41]. An SSVEP index of covert attention throughout the overt tracking of a moving target would allow for further clarification of how covert shifts of attention are influenced by target speed, and whether the effects pertain to the strength of covert shifts, the timing of such shifts, or both. Target-speed related modulation of covert peripheral attention is of particular concern in the domain of road-crossing safety, where increased vehicle speed may disproportionately affect individuals who tend to overtly track moving vehicles rather than covertly monitoring them through peripheral vision, as is the case with young children [42-43].

From a methodological perspective, the current study supports the use of the co-registration of eye-position recordings with SSVEP paradigms as a means investigating the dynamics of visual processing and attention while people perform tasks involving the tracking of moving objects. The development of this approach has recently shed light on the spread of attention during smooth-pursuit, with [24] providing electrophysiological evidence that visual attention is directed slightly ahead of targets as they move across the visual field, supporting behavioural results suggesting the same pattern [44-45]. A natural convergence of the current study with that of [24] would be to investigate the relationship between overt visual attention directed at a moving target and the default spread of attention while visual analysis of the target is taking place. The paradigm is also readily adaptable to investigate both overt and covert attention where multiple moving objects require selection or detection through either overt or covert visual attention [46]. The inclusion of a passive neural index of visual attention in such paradigms provides another layer of measurement when determining the timing or intensity or attentional shifts in complex visual environments. Methodologically speaking, there are a number of technical dimensions that must be addressed in order to obtain reliable SSVEP patterns that can be readily interpreted. The major concern is the control of low-level visual properties. It is imperative that participants’ eye-positions are monitored throughout the SSVEP trials, as the relative position of such stimuli in the visual field significantly modulates both the intensity and topography of the recorded signals [20, 47, 48]. This process will likely lead to the rejection of a certain number of trials involving inappropriate gaze-positions, and so the experimental planning needs to account for this reduction either by including a high number of trials, or an online index of gaze-accuracy which can repeat trials when necessary to compensate for rejected trials. In addition, it is likely that some tasks and conditions might involve differences in target-tracking accuracy, where specific conditions or contexts are more likely to elicit saccades that are difficult to inhibit (or in populations where such inhibition might be impaired). An analysis of trial rejection may therefore provide an index of this, as well as more in-depth analysis of gaze-behaviour in the trials as a means of relating such behavior with visual attention during periods of target pursuit [49]. However, experimental conditions with significantly different numbers of accepted trials might further complicate the interpretation of the comparison of SSVEP responses in these conditions as the signal-to-noise ratios in the EEG averages is strongly affected by this factor.

In summary, the application of SSVEPs to index overt visual attention while tracking a moving target provides a useful tool for understanding the effects of task-related covert attentional shifts in terms of both strength and timing. The results of the current study suggests that in contexts where covert attention is likely to be devoted to peripheral regions a concomitant reduction in overt attention to a moving target is likely to occur. The co-registration of EEG and eye-position while using the SSVEP technique would thus be well-suited to exploring this dynamic, and may provide valuable insight into areas where a reduction of overt attention due to covert distractors can lead to a decrease in performance or safety decisions relating to the moving targets.

## Supporting figures

**Fig 1.**
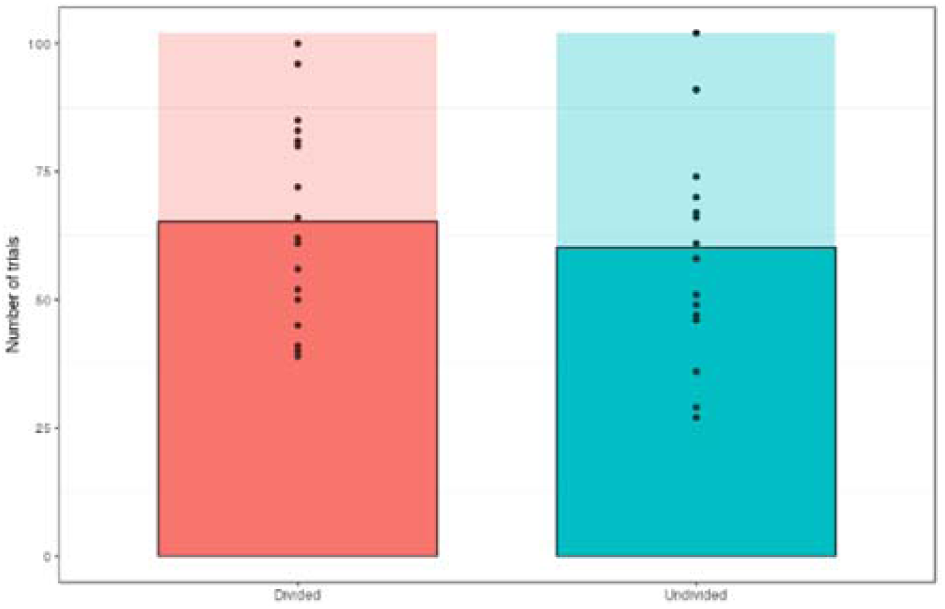
Trial numbers accepted in analyses. Trials were excluded due to unreliable tracking of the targets in both conditions from a total of 102 presented in each condition. Dots represent accepted trial numbers for each participant.

**Fig 2.**
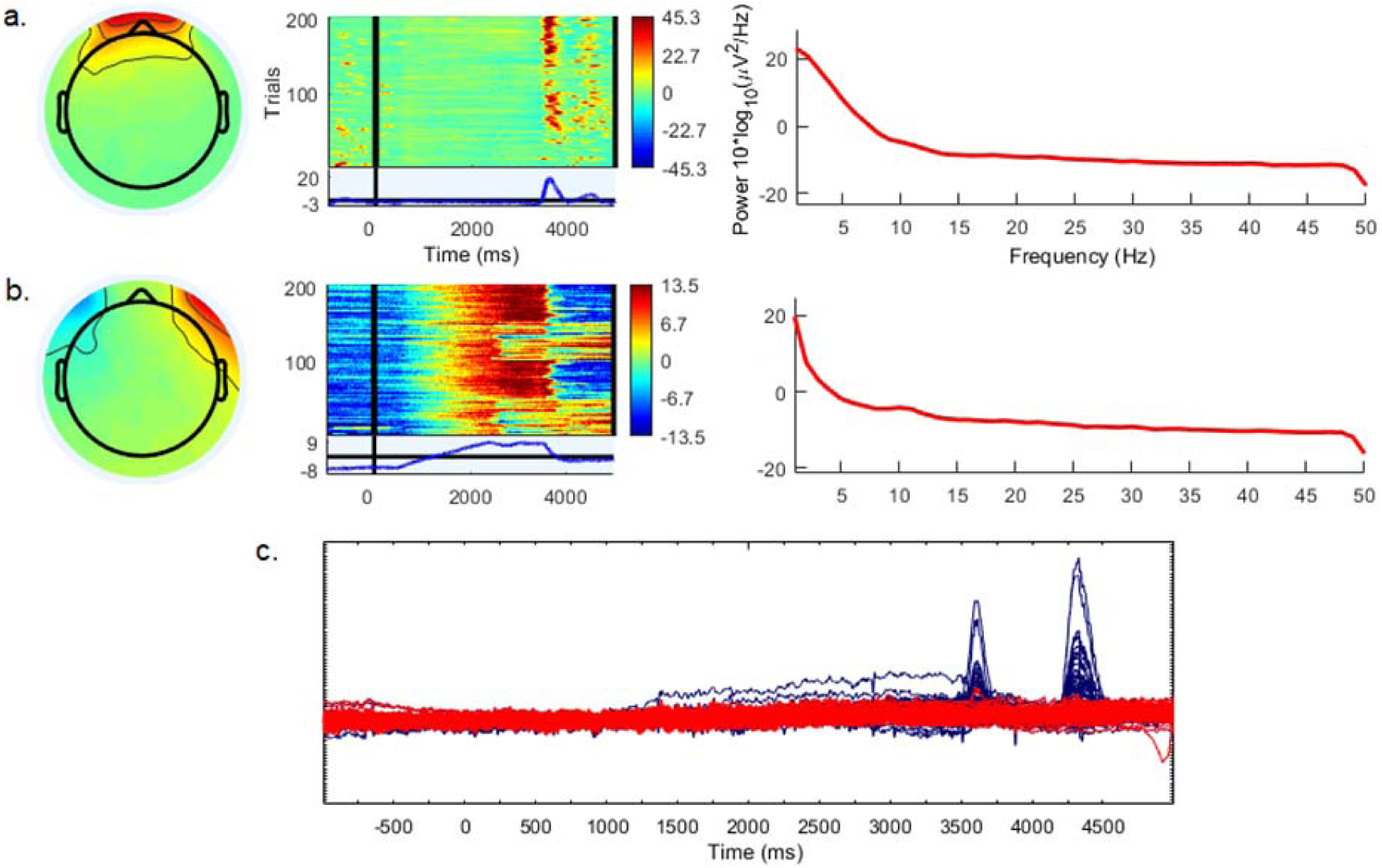
Examples of ocular distortion removal through independent component analysis. Distortions isolated and removed through the ICA process encompassed (a.) eye-blinks, and (b.) eye-movements involving saccades (sharp onset/offset activity), as well as smooth pursuit (slow drift). Fig 2c represents a single-trial example of uncorrected raw EEG (blue) and ICA corrected EEG (blinks, saccades, and drift removed) (red).

## Notes

### Competing Interest Statement

The authors have declared no competing interest.

### Summary of Updates

The manuscript text and figures have been revised for greater clarity and specificity.

## References

1. Kahneman, D. (1973). Attention and effort. Englewood Cliffs, NJ: Prentice Hall.

2. Lavie, N., Hirst, A., de Fockert, J.W., & Viding, E. (2004). Load Theory of Selective Attention and Cognitive Control. Journal of Experimental Psychology: General, 133(3), 339–354.

3. Lavie, N. (1995). Perceptual load as a necessary condition for selective attention. Journal of Experimental Psychology: Human Perception and Performance, 21, 451–468.

4. Lavie, N. (2005). Distracted and confused?: Selective attention under load. Trends in Cognitive Sciences, 9, 75–82.

5. Lavie, N. (2010). Attention, distraction, and cognitive control under load. Curr. Dir. Psychol. Sci. 19, 143–148.

6. Lavie, N., & Tsal Y. (1994). Perceptual load as a major determinant of the locus of selection in visual attention. Perception and Psychophysics, 56, 183–197

7. Moran, J., & Desimone, R. (1985). Selective attention gates visual processing in the extrastriate cortex. Science, 229, 782–784.

8. Reynolds, J. H., Chelazzi, L., & Desimone, R. (1999). Competitive mechanisms subserve attention in macaque Areas V2 and V4. J. Neurosci. 19, 1736–1753.

9. Sundberg, K.A., Mitchell, J.F., & Reynolds, J. H. (2009). Spatial attention modulates center-surround interactions in macaque visual area v4. Neuron, 61.

10. Posner, M.I. (1980). Orienting of attention. Quarterly Journal of Experimental Psychology, 32, 3–25.

11. Posner, M.I., Nissen, M.J., & Ogden, W.C. (1978). Attended and Unattended Processing Modes: The Role of Set for Spatial Location Editors’ Introduction.

12. Posner, M. I., Snyder, C. R., & Davidson, B. J. (1980). Attention and the detection of signals. Journal of Experimental Psychology: General, 109(2), 160–174.

13. Mishra, J., Zinni, M., Bavelier, D., & Hillyard, S.A. (2011). Neural basis of superior performance of action videogame players in an attention-demanding task. J. Neurosci. 31, 992–998.

14. Zhou, Y., Liang, L., Pan, Y., Qian, N., & Zhang, M. (2017). Sites of overt and covert attention define simultaneous spatial reference centers for visuomotor response. Sci. Rep. 7:46556.

15. Heinen S.J., Jin Z., & Watamaniuk S. N. (2011). Flexibility of foveal attention during ocular pursuit. Journal of Vision, 11, (2):9, 1–12.

16. Ludwig, C.J., Davies, J.R., & Eckstein, M.P. (2014). Foveal analysis and peripheral selection during active visual sampling. Proceedings of the National Academy of Sciences, 111:291–299.

17. Andersen S. K., & Muller M. M. (2010). Behavioral performance follows the time course of neural facilitation and suppression during cued shifts of feature-selective attention. Proceedings of the National Academy of Sciences, 107 (31), 13878–13882.

18. Kim, Y.J., Grabowecky M., Paller K. A., Muthu K., & Suzuki S. (2007). Attention induces synchronization-based response gain in steady-state visual evoked potentials. Nature Neuroscience, 10 (1), 117–125.

19. Morgan, S.T., Hansen, J.C., & Hillyard, S.A. (1996). Selective attention to stimulus location modulates the steady-state visual evoked potential. Proc Natl Acad Sci USA 93:4770–4774.

20. Müller, M.M., Teder-Sälejärvi, W., & Hillyard, S.A. (1998). The time course of cortical facilitation during cued shifts of spatial attention. Nat Neurosci 1: 631–634.

21. Störmer V. S., Winther G. N., Li S. C., & Andersen S. K. (2013). Sustained multifocal attentional enhancement of stimulus processing in early visual areas predicts tracking performance. Journal of Neuroscience, 33 (12), 5346–5351.

22. Toffanin P., de Jong R., Johnson A., & Martens S. (2009). Using frequency tagging to quantify attentional deployment in a visual divided attention task. International Journal of Psychophysiology, 72 (3), 289–298.

23. Norcia, A.M., Appelbaum, L.G., Ales, J.M., Cottereau, B.R., & Rossion, B. The steadystate visual evoked potential in vision research: A review. J Vision, 15:4.

24. Chen J., Valsecchi M., & Gegenfurtner, K. (2017a) Attention is allocated closely ahead of the target during smooth pursuit eye movements: evidence from EEG frequency tagging. Neuropsychologia, 102:206–216.

25. Chen, J., Valsecchi, M., & Gegenfurtner, K. R. (2017b). Enhanced brain responses to color during smooth-pursuit eye movements. Journal of neurophysiology, 118(2), 749–754.

26. Friedman, B. H., & Thayer, J. F. (1991). Facial muscle activity and EEG recordings: Redundancy analysis. Electroencephalography and Clinical Neurophysiology, 79, 358–360.

27. Goncharova, I. I., McFarland, D. J., Vaughan, T. M. & Wolpaw, J. R. (2003). EMG Contamination of EEG: Spectral and Topographical Characteristics. Clinical Neurophysiology, 114(9), 1580–1593.

28. Miskovic, V., & Keil, A. (2015). Reliability of event-related EEG functional connectivity during visual entrainment: magnitude squared coherence and phase synchrony estimates. Psychophysiology 52, 81–89.

29. Ales J. M., Farzin F., Rossion B., & Norcia A. M. (2012). An objective method for measuring face detection thresholds using the sweep steady-state visual evoked response. Journal of Vision, 12 (10): 18, 1–18.

30. Dienes, Z., (2011). Bayesian Versus Orthodox Statistics: Which Side Are You On? Perspectives on Psychological Science, 6, 274–290.

31. Wagenmakers, E.J., Love, J., Marsman, M. et al. (2018). Bayesian inference for psychology. Part II: Example applications with JASP. Psychonomic Bulletin & Review, 25(1), 58–76.

32. JASP Team (2019). JASP (Version 0.11.1)[Computer software].

33. Blair R.C., & Karniski W. (1993). An alternative method for significance testing of waveform difference potentials. Psychophysiology, 30:518–524.

34. de Brouwer, S., Yuksel, D., Blohm, G., Missal, M., & Lefevre, P. (2002). What triggers catch-up saccades during visual tracking? Journal of Neurophysiology, 87 (3), pp. 1646–1650.

35. Fehd, H.M., & Seiffert, A. (2010). Looking at the center of the targets helps multiple object tracking. Journal of Vision, 10: 1–13

36. Delorme, A., & Makeig, S. (2004) EEGLAB: an open source toolbox for analysis of single-trial EEG dynamics including independent component analysis. Journal of Neuroscience Methods, 134:9–21.

37. Vanegas, M. I., Blangero, A., & Kelly, S. P. (2015). Electrophysiological indices of surround suppression in humans. Journal of neurophysiology, 113(4), 1100–1109.

38. Seya, Y., & Mori, S. (2012). Spatial attention and reaction times during smooth pursuit eye movement. Attention, Perception, & Psychophysics, 74, 493–509.

39. Beck, D., & Lavie, N. (2005). Look here but ignore what you see: effects of distractors at fixation. Journal of Experimental Psychology, 31, 592–607.

40. Painter, D.R., Dux, P.E., Travis, S.L., & Mattingley JB. (2014). Neural responses to target features outside a search array are enhanced during conjunction but not uniquefeature search. J Neurosci. 34(9), 3390–401.

41. Bieg, H. J., Chuang, L. L., Bülthoff, H. H., & Bresciani, J. P. (2015). Asymmetric saccade reaction times to smooth pursuit. Experimental brain research, 233(9), 2527–38.

42. Biassoni, M., Confalonieri, F., & Ciceri, R. (2018). Visual exploration of pedestrian crossings by adults and children: Comparison of strategies. Transportation Research Part F, 56, 227–235.

43. Nicholls, V., Jean-Charles, G., Lao, J., de Lissa, P., Caldara, R. & Miellet, S. (2019). Developing attentional control in naturalistic dynamic road crossing situations. Scientific Reports, 9, 4176.

44. Khan, A.Z., Lefèvre, P., Heinen, S.J., & Blohm, G. (2010). The default allocation of attention is broadly ahead of smooth pursuit. J. Vision., 10(13):7, 1–17.

45. van Donkelaar, P., & Drew, A.S. (2002). The allocation of attention during smooth pursuit eye movements. Prog Brain Res., 140:267–277.

46. Lappin, J.S., Morse, D.L. & Seiffert, A.E. (2016). The channel capacity of visual awareness divided among multiple moving objects. Attention, Perception, & Psychophysics, 78: 2469.

47. Grgiĉ, R.G., Calore, E., & de’Sperati, C. (2016). Covert enaction at work: recording the continuous movements of visuospatial attention to visible or imagined targets by means of Steady-State Visual Evoked Potentials (SSVEPs). Cortex, 74, 31–52.

48. Punsawad Y, & Wongsawat Y (2017) A multi-command ssvep-based bci system based on single flickering frequency half-field steady-state visual stimulation. Med Biol Eng Comput, 55(6):965–977.

49. Renton, A., Painter, D., & Mattingley, J. (2018). Differential deployment of visual attention during interactive approach and avoidance behaviour. Cerebral Cortex, 29, 2366–2383.

